# TCRfinder: Improved TCR virtual screening for novel antigenic peptides with tailored language models

**DOI:** 10.1101/2024.06.27.601008

**Authors:** Yang Li, Chaoting Zhang, Xi Zhang, Yang Zhang

## Abstract

Accurate modeling of T-cell receptor (TCR) and peptide interactions is essential for immunoreaction elucidation and T-cell-based immunotherapeutic developments. We developed TCRfinder, a novel deep-learning architecture for TCR-peptide binding prediction and virtual screening. Large-scale benchmark experiments demonstrated a robust capability of TCRfinder in distinguishing interacting and non-interacting TCRs for unseen peptides, with accuracy significantly beyond current state-of-the-art methods. Furthermore, TCRfinder recognizes tumor neoantigen mutations from wild-type antigens of given TCRs, with a success rate nearly 50% higher than the best of existing methods. Detailed data analyses showed that the major advantage of TCRfinder lies in the specially trained TCR and peptide language models tailored with iterative attention network architecture, which can precisely reveal physical interaction patterns of cross-chain atoms and substantially enhance the precision of TCR-peptide interaction predictions. The open-source TCRfinder program can help facilitate large-scale deployment of high-quality TCR and neoantigen virtual screening, offering exciting potential for personalized TCR-based immunotherapies.

## INTRODUCTION

The T-cell receptor (TCR) is a protein complex found on the surface of T cells, playing a crucial role in the recognition of antigen peptides bound to the major histocompatibility complex (MHC) molecules. TCRs can recognize antigens, originating from viruses, bacteria, or somatically mutated genes (i.e., neoantigens), to activate cytotoxic T cells (CD8+ T cells) or helper T cells (CD4+ T cells), thereby cleaning up invading pathogens and tumor cells. As a result, the identification and characterization of TCRs that effectively interact with specific antigens are fundamental for understanding immune responses and designing targeted immunotherapies.

Previous studies primarily relied on experimental approaches, such as tetramer^1^, TetTCR-seq^2^ and T-scan^3^, to identify antigen-specific TCRs. However, these methods are not only time-consuming and costly but also have a general low success rate^4^. Thus, there is an urgent demand for effective TCR screening based on computational methods, which would significantly reduce the time and cost of identifying antigen specific TCRs.

The initial computational methods, including TCRdist^5^, DeepTCR^6^, TCRGP^7^, TCRex^8^, and NetTCR^9^, utilized various approaches such as CDR similarity-weighted distances, random forests^10^, Gaussian process classification methods^11^, and convolutional neural networks (CNNs) to distinguish between positive and negative TCRs for antigenic peptides. However, their utility is constrained, particularly when involving novel or unseen peptides that have not been encountered in the training datasets. Recent advancements in TCR screening methods aim to overcome the limitation of applicability to unseen peptides in methodology. For example, pMTnet^12^ employs deep transfer learning and leverages pretrained autoencoders, to predict interactions between TCRs and peptides. ERGO^13,14^ and DLpTCR^15^ utilize CNNs, or ensembles of CNNs, to recognize TCR–antigen binding. More recent approaches, such as ATM-TCR^16^, AttnTAP^17^, and PanPep^18^, incorporate deep learning models with attention mechanisms to predict these interactions. While these methods demonstrate the potential to identify TCR–peptide interactions for unseen peptides, their accuracy remains suboptimal^19^, likely due to the constraints imposed by the limited availability of data.

In this work, we introduce TCRfinder, a novel approach that harnesses the power of language models^20^ (LMs) to address the challenge posed by limited data in virtual TCR and antigen screenings. Specifically, we have trained two special language models, i.e., TCR LM and peptide LM, from scratch to enhance the representations of TCR and peptide sequences. We found that the two LMs exhibit an improved prediction power for different TCR-peptide interactions and achieve the best overall performance, when integrated with an iterative attention network architecture. TCRfinder represents a significant stride toward more reliable TCR and antigen virtual screening, showcasing the potential of language models in advancing our understanding of immune system responses to diverse antigens.

## RESULTS

The core of the TCRfinder is a deep learning model trained for accurately scoring the interactions between the antigenic peptides and TCR sequences. Because the CDR3 region of TCR β-chain is the key determinant of the interactions, our model is trained mainly on the β-chain CDR3 regions of the TCR sequences. As outlined in **Figure 1A**, TCRfinder first employs a Joint Embedder module (**Figure 1D**), which utilizes two pre-trained LMs (**Figure 1B**) for peptide and TCRβ CDR3 sequences separately, followed by iterative transformer block learnings starting with concatenated and split embeddings (**Figure 1C**). The final TCR-peptide interaction scores are derived through a multi-layer perceptron (MLP) network based on the concatenated representations of TCR and peptide embeddings from the preceding transformer networks (**Figure 1**).

**Figure 1.**
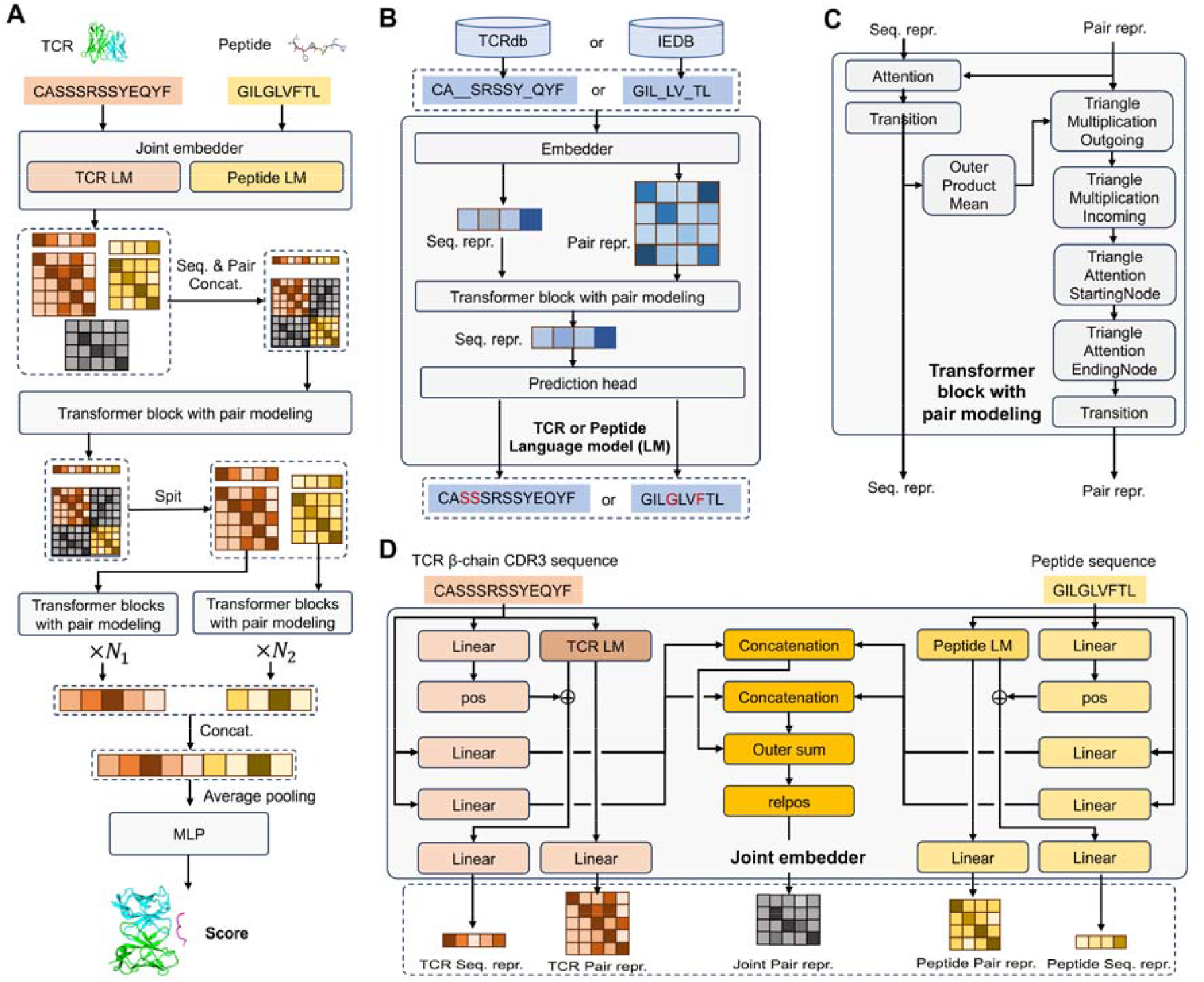
Flowchart of TCRfinder for sequence based TCR and peptide interaction predictions using deep learning networks. (A) Overview of the TCRfinder pipeline. (B) Detailed architecture of TCR and peptide language models. (C) Details of transformer block with pair modeling, utilized in both TCRfinder model and TCR and peptide language models. (D) Detailed architecture of Joint Embedder module to encode TCR and peptide sequences into sequential and pair representations.

### Specifically trained language models enhance specificity of sequence representations

Machine learning-based TCR virtual screening for unseen peptides is challenging because, by definition, the interaction data between TCRs and the unseen peptides is limited. To address the limitation, we trained two distinct language models tailored for TCR (CDR3 region of the β chain) and peptide sequences, with the aim to incorporate additional TCR and peptide sequence representations to better model the inherent relationships between these sequences. It is worth noting that there are already large language models designed for general protein sequences, e.g., ESM21,22. However, we posited that TCR and peptide sequences exhibit unique sequence patterns that may need specific model training.

In **Figure 2A**, we show a head-to-head comparison of perplexities of the ESM-2 model (t33_650M_UR50D) and the TCR LM trained from scratch by TCRfinder on 884 validation TCR sequences from VDJdb^23^ that have a maximum sequence identity of 80% to the training TCRs (see **Text S1** in **Supporting Information, SI**). Here, the (pseudo) perplexity of a LM for a protein sequence is computed by

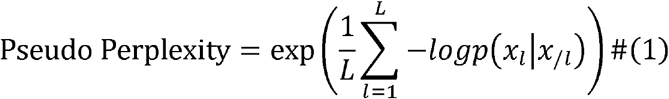

where *L* is the sequence length, *x*_*l*_ the masked residue at position *l*, and *x*_/*l*_ the context. The value of the perplexity can serve as a metric to quantify the mean uncertainty of the LM, reflecting how well the LM is able to predict the masked token in the sequence. A lower perplexity indicates that the model is more certain and accurate in its predictions, approaching the ideal scenario of one. In contrast, a higher perplexity suggests higher uncertainty and a less effective model, with values approaching the number of unique tokens, indicating randomness in predictions. Please note that for the evaluation of the TCR LM, we specifically computed the perplexity of the 7 center amino acids of the CDR3 region, as these central regions are known to be more flexible.

**Figure 2.**
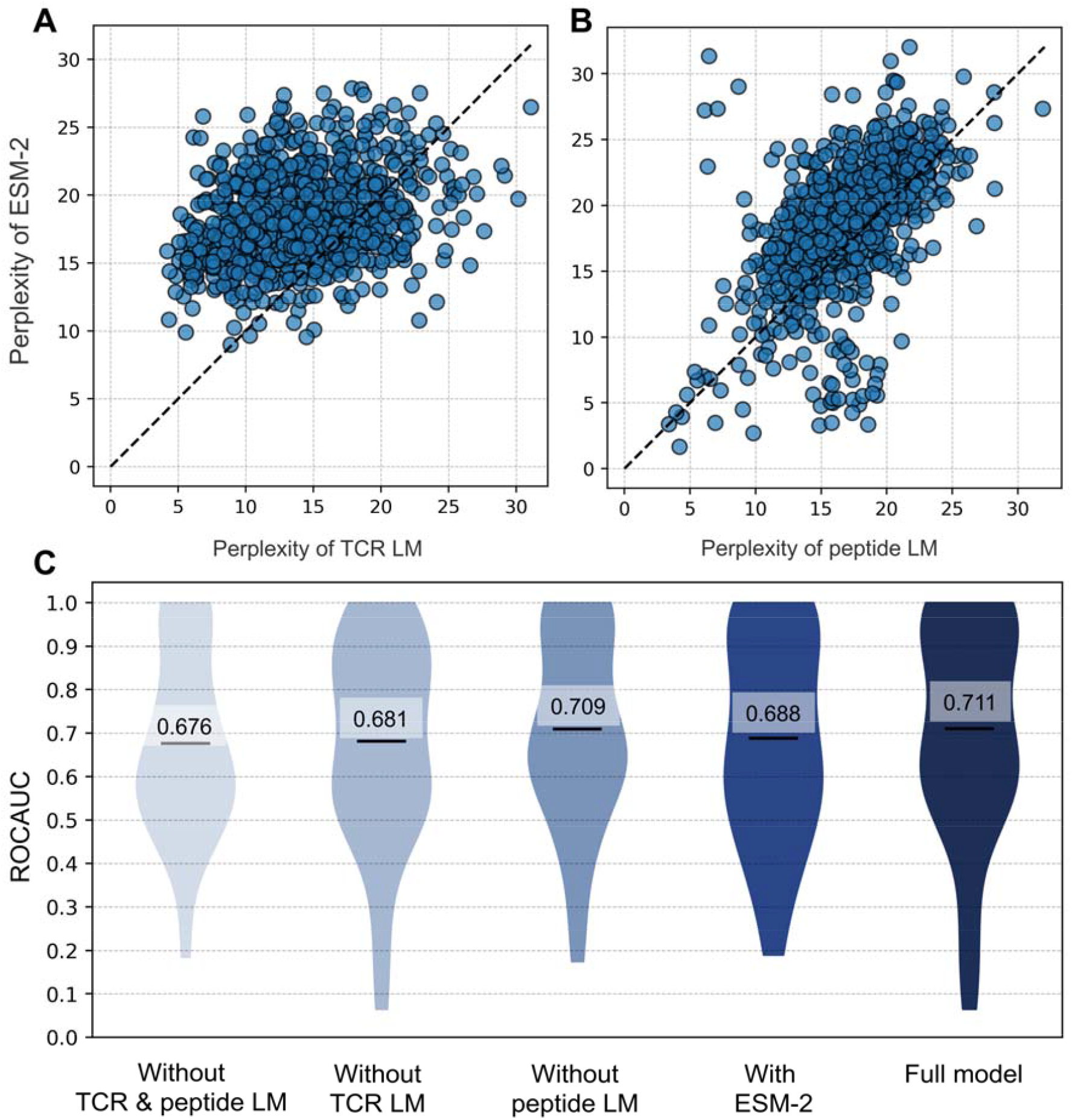
Comparative Analysis of Language Models in TCRfinder. (A) Head-to-head perplexity comparison of ESM-2 and TCR LM models on TCR β-chain CDR3 sequences. Head-to-head perplexity comparison of ESM-2 and specially trained peptide LM models on peptide sequences. (C) ROCAUC of TCR screening experiments on the validation dataset with and without using language models.

It is shown that the TCR LM trained in TCRfinder achieves an average perplexity of 14.289, significantly lower than the 18.321 by ESM-2, with a P-value of 4.98E-100 in paired one-sided Student’s t-test. Out of the 884 test cases, 78.96% (698 cases) exhibited a lower perplexity with TCR LM, compared to only 21.04% (186 cases) for ESM-2. The superiority of the TCR LM over ESM-2 is understandable because ESM-2 has been trained on general protein sequences and is therefore less sensitive to the specific patterns present in TCR sequences. These results also indicate that TCR sequence patterns significantly diverge from general protein sequence patterns, highlighting the need for a customized strategy when dealing with the specific patterns of TCR sequences.

In **Figure 2B**, we also present a perplexity comparison of ESM-2 and the peptide LM trained in TCRfinder on 1000 validation peptides (see **Text S1**), where the specially trained peptide LM demonstrates again a significantly lower perplexity (17.055) than ESM-2 (18.758) with a P-value of 6.54E-40 in a one-sided Student’s t-test. Furthermore, the peptide LM achieves a lower perplexity in 755 out of 1000 cases (75.50%), suggesting its higher efficacy in capturing peptide sequence patterns compared to ESM-2.

### Impacts of language models on peptide specific TCR virtual screening

One primary goal of training TCR and peptide LMs is to enhance the modeling accuracy of TCR and peptide associations. To examine the impacts of these specially trained LMs on TCR virtual screening, we present in **Figure 2C** a comparison of ROCAUC achieved by TCRfinder with and without using LMs on 58 peptides randomly selected from the validation dataset (**Text S2**). Here, ROCAUC stands for ‘Areas under the Receiver Operating Characteristic curve’ computed in the classification experiment distinguishing interacting from non-interacting TCRs for the given antigenic peptides. For the case without LMs, TCRfinder models are trained with LM matrices replaced by the query sequences only in the Joint Embedder module shown in **Figure 1D**.

It is shown that the incorporation of LMs significantly improves the screening performance, with the average ROCAUC of full TCRfinder (0.711) being 5.2% higher than that without using LMs (0.676). The ablation experiment shows that the major impact comes from the TCR LM, as the ROCAUC of TCRfinder after removing TCR LM is 4.4% lower than the full TCRfinder while removing peptide LM reduces ROCAUC only by 0.3%. The varied impact of LMs on the performance of TCR virtual screening might stem from the lower specificity of the peptide language models, especially for the unseen peptides in this validation dataset. This result is consistent with the data in **Figures 2A** and **2B**, where the perplexity of the TCR LM (14.289) is considerably lower than that of the peptide LM (17.055), indicating better sequence pattern recognition power of the TCR over peptide LM.

In **Figure 2C**, we also list the average ROCAUC value (0.688) of TCRfinder when replacing specially trained LMs by ESM-2 models, which is considerably lower than that of original TCRfinder models (0.711). This result demonstrates again the importance of using specially trained LMs over general protein LMs, even though the latter has been trained in a much larger sequence space with significantly higher dimension of models^21,22^ (8.7 million short-length vs ∼50 million full-length sequences with 6.1 million vs 651 million parameters for the TCRfinder and ESM-2 LMs, respectively).

### TCRfinder outperforms previous methods for TCR virtual screening

To systematically examine the ability of TCRfinder in TCR virtual screening, we collected a set of 38 nonredundant peptides from the VDJdb, each with 1-239 interacting TCRβ CDR3 sequences unseen in any of the TCRfinder training datasets (see **Text S2**). **Figure 3** summarizes TCR virtual screening results of TCRfinder, in control with six state-of-the-art third-party methods, including ATM-TCR^16^, PanPep^18^, ERGO^13,14^, AttnTAP^17^, pMTnet^12^ and DLpTCR^15^, which are all installed in our local computers and implemented with default parameters.

**Figure 3.**
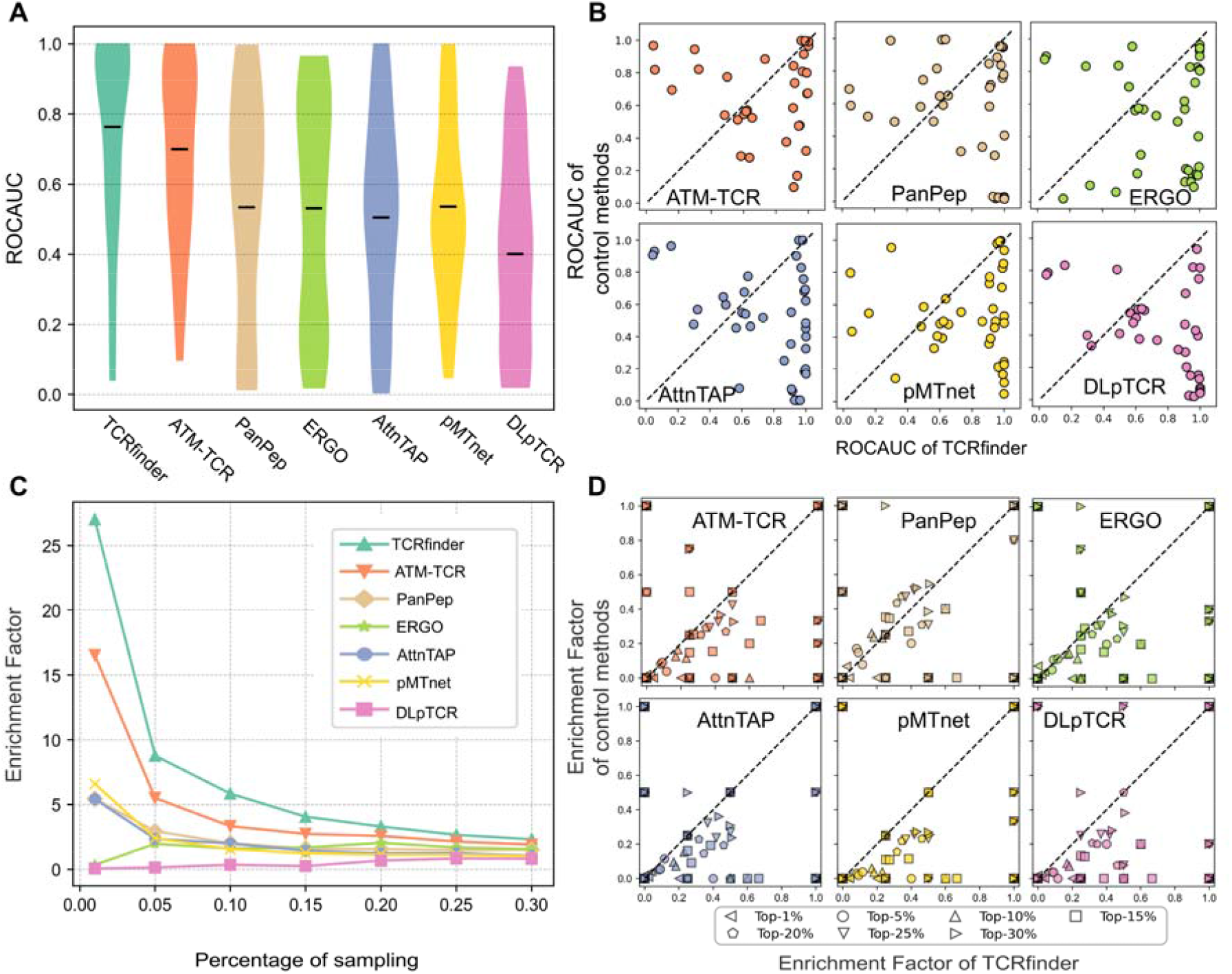
The comparison of TCRfinder with the control methods in TCR screening on 38 unseen peptides. (A) Distribution of ROCAUC with the central mark indicating the mean. (B) Head-to-head ROCAUC comparison. (C) Enrichment Factors (EFs) versus percentage of sampling. (D) Head-to-head comparisons of EFs that are scaled to [0.0, 1.0] by the theoretical maximum EF value at corresponding top-*x*% cut-offs.

In **Figure 3A**, it is observed that most methods exhibit limitations in effective TCR recognitions for unseen peptides, where five control methods (PanPep, ERGO, AttnTAP, pMTnet, and DLpTCR) generated average ROCAUC values of 0.534, 0.531, 0.504, 0.536, and 0.401, respectively, marginally surpassing or even falling below a random guess (ROCAUC = 0.5). Although ATM-TCR produces a considerably higher ROCAUC= 0.699 than other control methods, TCRfinder achieved the highest ROCAUC of 0.763, marking a 9.2% increase over ATM-TCR. **Figure 3B** provides a detailed head-to-head comparison of peptide-wise ROCAUC between TCRfinder and control methods, where TCRfinder achieves a higher ROCAUC in 28 cases compared to ATM-TCR, whereas ATM-TCR accomplishes this in 10 cases. The numbers are 27/11, 32/6, 27/11, 29/9, and 32/6, compared to PanPep, ERGO, AttnTAP, pMTnet and DLpTCR, respectively.

Notably, TCRfinder has a median ROCAUC of 0.912, with majority of test peptides scoring between 0.8 and 1.0, representing a noteworthy 20.5% improvement over ATM-TCR that has a median ROCAUC of 0.757. The superiority of TCRfinder in median ROCAUC is particularly meaningful to experimentalists, as such a high median ROCAUC suggests that with TCRfinder screening models, for half of the unseen peptides, experimentalists only need to validate, on average, less than 10% (equals to 1-0.912) of the sample pools to identify the true interacting TCRs. The result highlights the practical usefulness of effective computational screening in helping significantly reduce the time and cost of tumor specific TCRs recognitions and immunotherapies.

In **Figure 3C**, we extend our analysis to the comparisons of Enrichment Factors (EFs) which is defined as

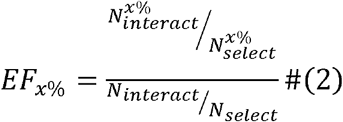

where *N*_*interact*_ and *N*_*select*_ are the total number of interacting and all TCRs in the screening pool, respectively. 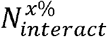 and 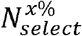 are the numbers of true interacting TCRs and the number of all candidates in the top-*x*% of the TCRs selected on the scores predicted by the corresponding methods. Generally, a higher EF value indicates better performance and *EF* = 1 indicates random selection. In this work, a wide range of top-*x*% cut-offs (i.e., 1%, 5%, 10%, 15%, 20%, 25% and 30%) are used to compare the performance under different scenarios.

Consistent with the ROCAUC results, ATM-TCR produces consistently higher EFs than other control methods, where the P-values between TCRfinder and ATM-TCR, PanPep, ERGO, AttnTAP, pMTnet and DLpTCR are 4.54E-03, 2.25E-05, 9.46E-08, 9.01E-06, 7.87E-06 and 3.32E-10, respectively, across all the top-*x*% cut-offs. For the top-1% sample cutoff, for instance, TCRfinder achieves an average EF=27.01, which is 63.2% higher than ATM-TCR (16.53) and >4 times higher than that of the other three methods (PanPep, AttnTAP, and pMTnet) that have achieved better than random EF (i.e., EF > 1.0).

**Figure 3D** further lists head-to-head comparisons of EF across multiple cut-offs between TCRfinder and control methods. Here, EF values are scaled to [0.0, 1.0] by the theoretical maximum value of the corresponding top-*x*% cut-offs, i.e., 100/*x*, for improved visualization. Out of 266 indices (i.e., 38 peptides with 7 top-*x*% cut-offs each), TCRfinder consistently achieves higher or equal scaled EF in 238, 224, 238, 235, 256, and 255 cases compared to ATM-TCR, PanPep, ERGO, AttnTAP, pMTnet, and DLpTCR, respectively. In contrast, the control methods generate better or equal performance in at most 172 cases. The associated P-values, with a maximum of 9.20E-07, for the scaled EF, further validate the statistical significance of TCRfinder’s superior performance across a spectrum of enrichment cut-offs.

### Case studies reveal effectiveness of unsupervised learning for cross-chain structure interactions

To further examine the strength and weakness of TCRfinder, as case studies we look into details of two peptides of ‘**MMWDRGLGMM’** and ‘**EVDPIGHLY’**, which represent two typical examples standing at the upper-left and bottom-right corners of ROCAUC-homology plot **Figure S1**.

First, ‘**MMWDRGLGMM’** is a synthetic epitope presented by HLA-A*02^24,25^ with only one interacting TCR identified in VDJdb. The CDR3 sequence ‘*CASSLSFGTEAFF*’ of this TCR is composed of the V and J genes of TRBV6-4 and TRBJ1-1, respectively. As shown in **Figure S2A**, the most similar peptide in the training dataset is ‘NMMWFQGQL’, which is another synthetic epitope presented by the same MHC, with a sequence identity of 40% to the test peptide. However, the two known interacting TCRs for ‘NMMWFQGQL’, with β-chain CDR3 sequences of ‘*CASSRDTVNTEAFF*’ and ‘*CASSRDFVSNEQYF*’, have completely different J genes (TRBJ1-1 and TRBJ2-7). Thus, the high ROCAUC of 0.985 (Figure **S1)** is not due to the simple homologous transfer from the training dataset.

The TCR-peptide interacting complex structure from PDB (PDBID: 6AMU) is depicted in **Figure 4A**, with a detailed inter-chain Cα distance map presented in **Figure 4B**. Interestingly, the central region of the TCR-peptide distance map, which corresponds to physical inter-residue interactions, shows a similar pattern to the mean attention map which was automatically learned from solely TCR-peptide sequence data by TCRfinder. Here, the attention map between the TCRβ CDR3 region and peptide sequences was extracted directly from the transformer block following the Joint Embedder and represents the mean of attention weights from each head. Additionally, focusing on the TCRβ CDR3 region, **Figure 4C** illustrates the minimum distance between each residue of TCRβ CDR3 and the peptide residues, superposed with the accumulated attention weights corresponding to each residue of TCRβ CDR3. A significant correlation (PCC of 0.549) can be observed between the two signals, despite no structural information being included during the TCRfinder training. These findings suggest that the attention mechanism in TCRfinder captures meaningful information regarding the spatial binding interface of residues in the TCRβ CDR3 region and their interactions with the peptide, an ability critical to the enhanced TCR-peptide interaction predictions.

**Figure 4.**
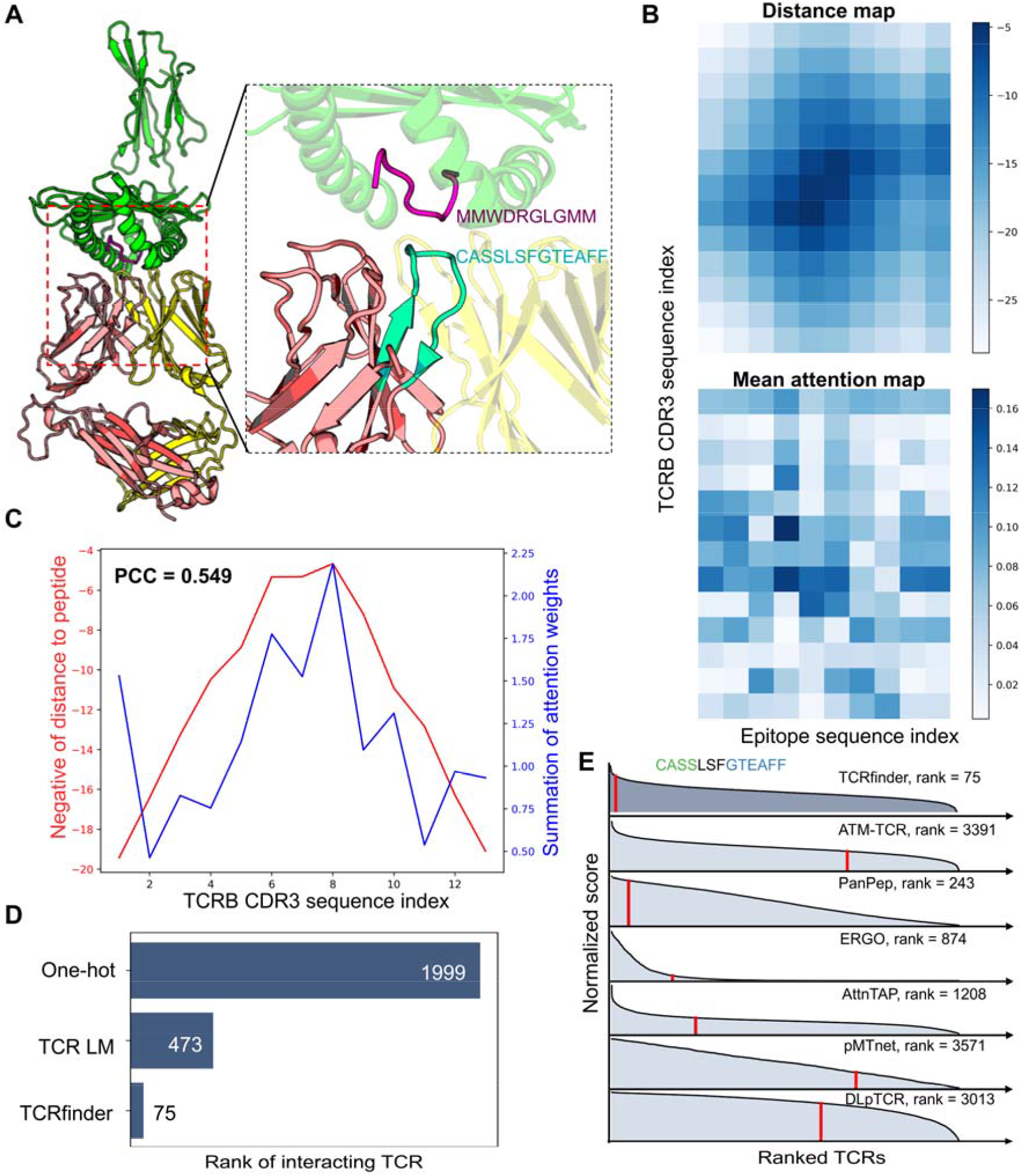
Case study of TCR screening for peptide ‘MMWDRGLGMM’. (A) Experimental structure of TCR-peptide interacting complex with TCRβ CDR3 region and the peptide highlighted in cyan and purple, respectively. (B) Distance map between interchain Cα atoms, attention map between TCRβ CDR3 region and peptide sequences.(C) Residue wise minimal distance and attention weights to the peptide. (D) Comparison of TCR LM encoding and one-hot encoding in a nearest-neighbor approach for the studied case. (E) identification of the interacting TCR for the case peptide by TCRfinder and other methods.

To further evaluate the effectiveness of TCR LM in this context, a basic nearest-neighbor approach is implemented by encoding TCRs using both TCR LM and the basic one-hot encoding. Candidate TCRs are searched by comparing the representations of interacting TCRs of the template peptide in VDJdb and ranked according to the distance of representative vectors between two TCRs. In this approach, the residue-wise representations of TCRs can be simply averaged along the sequence axis into a fixed-length vector. The ranks of interacting TCR for TCR LM encoding and the basic one-hot encoding are 473 and 1999, respectively, as shown in **Figure 4D**. This demonstrates that TCR LM learns more informative representations than just residue types. Leveraging TCR LM, along with further supervised training, TCRfinder successfully identifies the interacting TCR at the 75^th^ trial from a pool of 5000 background TCRs, constituting only 1/3 of the second-best ranking achieved by PanPep (243^rd^) (**Figure 4E**). In contrast, other control methods exhibit rankings exceeding 800 and, in some cases, > 3500. These results strongly affirm the robust capability of the proposed TCRfinder to effectively screen interacting TCRs, not only for unseen peptides but also for peptides that are distinctly novel and challenging.

In the second example of ‘**EVDPIGHLY**’, TCRfinder achieved a poor ROCAUC of 0.042, even though a highly homologous peptide (MEVDPIGHLY) exists in the training dataset with 100% sequence identity (or 90% if normalized by the length of ‘MEVDPIGHLY’), and both peptides belong to the MAGE-A3 epitope.

As shown in **Figure S2B**, the interacting TCR for the training peptide has a β-chain CDR3 sequence of ‘*CSANPRTTLYEQYF*’, formed by V and J genes of TRBV20-1 and TRBJ2-7, respectively. In contrast, the interacting TCR for the test peptide has a distinct β-chain CDR3 sequence of ‘*CASSFNMATGQYF*’, with V and J genes of TRBV5-1 and TRBJ2-7, respectively. The notable differences in TCRs despite the similar peptide sequence make the virtual screening challenging. Furthermore, 84.14% of the randomly selected background TCRs exhibited a higher sequence similarity to the positive TCR in the training set. Consequently, TCRfinder failed to achieve a reasonable ranking for the interacting TCR.

Notably, in VDJdb, the test and training peptides are presented by different MHCs, i.e., HLA-A*01 and HLA-B*18, respectively. As data accumulates, we aim to incorporate more metadata (e.g., species and MHC types) and peripheral protein environments into consideration for such cases when they are available.

Despite these challenges, it’s noteworthy that for 8 peptides with a maximum sequence identity exceeding 80%, 6 (75%) of them demonstrated high ROCAUC values of at least 0.906 and an average rank of 202.23 out of 5000 candidates by TCRfinder (**Figure S1**).

### Recognition of neoantigens against wildtype peptides

Neoantigens are newly formed antigen peptides generated by mutations in the DNA of cancer cells. A critical step in T-cell based cancer therapeutics is the identification of TCRs that can recognize neoantigens presented by MHC proteins^4,26^. However, since the difference between a neoantigen and corresponding wild-type peptide often involves only a single residue mutation, neoantigen-reactive TCRs might also interact the corresponding wild-type peptide. Such interactions could lead to serious autoimmune disorders, creating an urgent need to identify neoantigen-reactive TCRs that are less likely to interact with wildtype peptides. But the high sequence similarity between between neoantigen and wildtype peptides often renders the computational screening of neoantigens for given TCR highly challenging.

The default TCRfinder model cannot be directly applied to neoantigen screening, as the loss function focuses solely on the ranking between two TCRs for a given peptide (see **Eq. 5** in **METHODS**). To address this, we developed an additional model specifically focused on scoring interactions between diverse peptides for a given TCR as described in ‘**Alternative model trained for TCR-based neoantigen screening**’ in **METHODS**. To quantitatively assess the capability of the model, we gathered a total of 64 independent TCR-peptide pairs, including 32 experimentally validated TCR-neoantigen interactions and 32 corresponding TCR-wildtype non-interacting pairs, also validated experimentally^27-29^. Here, we used the data from VDJdb for training, where all pairs sharing the same TCRs or peptides to the 64 test TCR-peptide pairs have been excluded from the training set to prevent data contamination (**Text S2**).

In **Figure 5A**, we present a summary of success rate of TCRfinder on the 64 TCR-peptide pairs in control with other six third-party programs, where a success case refers to that in which the program correctly prioritizes the TCR-neoantigen interaction with a higher score than the TCR-wildtype peptide pairs. As shown in **Figure S3**, all the control methods demonstrate relatively low success rates ranging from 0.375 (ERGO) to 0.563 (DLpTCR), indicating the challenging nature of this task. In contrast, TCRfinder achieved a remarkable success rate of 0.844, due to the special training strategy. As shown in **Figure 5B**, the P-value between the binding scores with neoantigen and wildtype peptides by TCRfinder is 1.34E-07 in paired one-sided Student’s t-test (**Figure 5B**), highlighting its robustness in distinguishing TCRs with distinct interactions between neoantigens and wild-type peptides.

**Figure 5.**
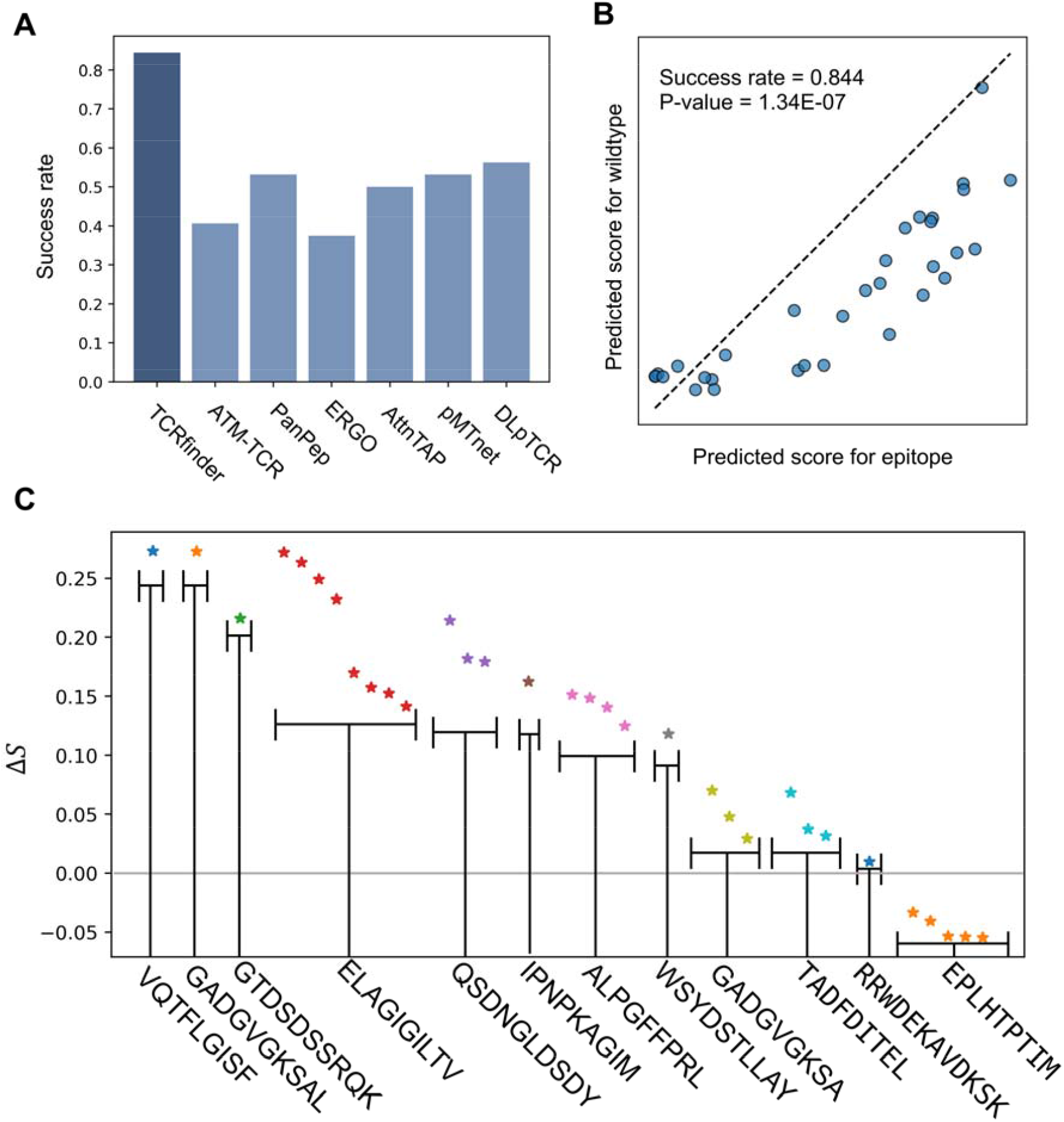
Test results of neoantigen recognition on 64 TCR-peptide pairs. (A) Success rate of different programs in correctly recognizing TCR-neoantigen interactions. (B) Predicted interaction score for TCR-neoantigen versus that for TCR-wildtype peptide by TCRfinder. (C) Score difference between TCR-neoantigen and TCR-wildtype pairs by TCRfinder as grouped by distinct neoantigens.

The 32 TCR-neoantigen interactions involve a total of 12 unique neoantigens. **Figure 5C** lists the details of score differences, Δ*S* = *S*_*neoantigen*_ – *S*_*wildtype−peptide*_, on the interactions grouped by neoantigens. It can be observed that TCRfinder correctly recognizes 11 out of 12 (91.67%) neoantigens with positive Δ*S*, failing only in one neoantigen, ‘EPLHTPTIM’. Such a high success rate at the neoantigen level is critical for guiding the selection of TCRs with lower “toxicity” for personalized immunotherapies.

## DISCUSSION

Accurate modeling of TCR and antigenic peptide interactions is essential for elucidating immune responses and developing T-cell based immunotherapies. By harnessing specifically trained TCR and peptide language models, we introduced TCRfinder, a novel deep learning method for precise TCR-peptide binding interaction prediction. Through a careful examination of various performance metrics, we found the robust capability of TCRfinder in distinguishing interacting and non-interacting TCRs for unseen peptides, with accuracy significantly beyond current state-of-the-art methods. Furthermore, through rigorous training and validation, TCRfinder demonstrated a remarkable ability to distinguish TCRs with distinct interactions between neoantigens and wild-type peptides, with a success rate nearly 50% higher than the best existing method. This capability is particularly important to minimize toxic autoimmune responses in T-cell based immunotherapies.

Several unique designs have contributed to the superior performance of the TCRfinder. First, the specifically trained TCR and peptide language models exhibit higher specificity (or lower perplexity) than general large protein language models, such as ESM-2, allowing better modeling of evolutionary patterns in diverse TCRs (particularly the sensitive β-chain CDR3 regions) and peptide sequences. Second, as demonstrated by the case studies, the specially designed Joint Embedder network, when integrated with the LMs, can reveal meaningful information regarding the spatial binding interface of residues within the TCRβ CDR3 region, highlighting its potential to bridge the gap between sequence-based predictions and structural insights. Third, although both TCR and neoantigen screens rely on the specific TCR-peptide interactions, the procedures involve different sampling backgrounds (i.e., one on multiple TCRs for a given peptide and another on highly similar peptide sequence for a given TCR). Therefore, separate models trained with specially designed loss functions can further improve the accuracy and specificity of the screening experiments.

Nevertheless, there are still substantial limitations in the TCRfinder model. Frist, despite careful training, the peptide LM is still much less specific than the TCR β-chain CDR3 region LM and thus has a less significant impact on TCRfinder’s overall performance. One reason could be the relatively small training dataset, where the trained LM cannot accurately capture the diversity of TCR-bound peptide sequence patterns. Second, the current TCRfinder model has difficulty in recognizing the closely homologous peptides that bind with distinct TCRs. This limitation may also relate to the limited sensitivity of the specially trained peptide LMs. Collecting larger datasets of peptide/neoantigen sequences and incorporating peripheral features such as HMC and TCR sequences in the LM training might help address the issues.

In summary, our findings highlight the significance of leveraging language models in TCR virtual screening, offering a powerful approach to predict antigen-specific TCRs for unseen antigenic peptide. Such advancement holds promise for applications in individualized TCR-related therapies.

## METHODS

TCRfinder is a deep learning pipeline designed for sequence based TCR and neoantigen screenings. The core strategy of the pipeline is the development of accurate interaction scoring models from the sequences of the β-chain TCR CDR3 region and target peptides, which consists of four steps (**Figure 1A**). Firstly, two language models (TCR LM and peptide LM) are pre-trained for the TCR and peptide sequences separately. A joint embedder model is then learned from the concatenated representations of the TCR and peptide LMs. Next, transformer blocks are implemented iteratively on both paired and individual TCR and peptide representations to create new TCR and peptide embedding matrixes. Finally, a TCR-peptide interaction scoring model is obtained from a Multi-Layer Perceptron (MLP) block layer based on the new TCR and peptide embeddings. To address issues of TCR and neoantigen recognitions, two models on peptide-based TCR screening and TCR-based neoantigen screening are trained separately.

### TCR and peptide language model training

The neural network architecture of TCR and peptide LMs is presented in **Figure 1B**. Given a masked TCR sequence, the embedder layer is employed to extract hidden representations in the forms of both sequential and pair matrices. The detailed flowchart of the Embedder Module is shown in **Figure S4**, where the input is the one-hot encoding of the masked TCR or peptide sequence, and the resulting output contains sequential and pair representations. Specifically, the sequential representation is derived through a linear transformation over the input features, added with 1-D positional encoding. The pair representation is the outer sum of two separate hidden features, each obtained by individual linear transformations. Additionally, the pair representation is enriched by the inclusion of 2-D positional encoding.

Next, the sequential and pair representations serve as inputs of a set of transformer blocks, designed to model interactions between residues in the sequence (see **Figure 1C** as well as the description in ‘**Transformer Block with pair modeling**’ below). While the transformer block generates both sequential and pair representations, we only utilize the last sequential representation for predicting the residue type of the masked regions.

To train the TCR LM, we collected all downloadable TCR sequences from TCRdb^30^, comprising a total of 7,259,306 TCR β-chain CDR3 sequences, while the validation set for the TCR LM contains 884 TCRs included in VDJdb^23^, which have the maximum sequence identity of 80% to the training TCRs. In the case of the peptide LM, we collected all unique linear peptides from IEDB^31^ and randomly selected 1000 peptides for evaluation, with the remainder allocated for training, where a maximum sequence identity cutoff of 80% is maintained between the evaluation and training sets for peptide LM (see **Text S1** in **SI**).

For both TCR and peptide LMs, we set the number of blocks as 4 and 7, respectively. The dimensions for sequential and pair representations in the TCR LM are set as 128 and 96, while for the peptide LM, they are set as 256 and 96, respectively. Two masking policies are employed during training: (1) Randomly masking 10% of the residues; (2) Randomly masking a continuous region with a length of approximately 10% of the sequence length.

Both TCR and peptide LMs were trained using the Adam optimizer^32^ with default parameters for approximately 10 epochs. The training process was guided by minimizing the cross-entropy loss of the masked regions given a sequence, as defined by:

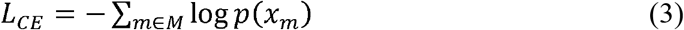

where *p*(*x*_*m*_)is the softmax probability for the *m*th masked residue and *M* refers to the masked residue set.

### Joint embedding of TCR and peptide sequences

Built on the pre-trained LMs, TCRfinder employs the Joint Embedder module to transform inputs into diverse representations. As illustrated in **Figure 1D**, the TCR and peptide sequences are initially encoded using one-hot encoding. Each sequence undergoes a linear transformation, accompanied by 1-D positional encoding injection (denoted as ‘pos’ in the architecture), resulting in hidden sequential representations. Simultaneously, the one-hot encoding features are input into their respective LMs to derive sequential and pair representations for TCR and peptide, respectively. Note that the representations from LMs will be projected by the linear layers to the desired dimensions. The output sequential representations are obtained by element-wise additions of (1) the linear layer with positional encoding and (2) the corresponding language model’s output after projection. The pair representations after the projection will also be considered as the outputs of the Joint Embedder.

On the other hand, the sequential representations of TCR and peptide will be fed into two linear layers individually. The resultant sequential representations are concatenated separately to create two joint sequential representations. A subsequent Outer Sum operation is applied to these joint sequential representations, resulting in the joint pair representation. The joint pair representation will have the 2-D positional encoded (denoted as ‘relpos’ in the **Figure 1D** architecture). The dimension sizes for sequential and pair representation are uniformly set to 64 and 48, respectively.

### Transformer Block with pair modeling

The joint sequence and pair representations from the Joint Embedder will be considered as the input of a single transformer block to capture interaction patterns. This simplified design aligns with the philosophy that a compact deep learning model can effectively learn intricate interaction patterns when transferring strong prior information from, for example, pretrained language models. The architecture of the transformer block is an extension inspired by the Evoformer in AlphaFold2^33^, as detailed in **Figure 1C**.

The training process begins with sequential and pair representations being through a sequence row-wise gated self-attention layer. Notably, the pair representation is transformed and added as the bias term of the attention map, facilitating the effective incorporation of relationships between residues in the output sequential representation. A sequence transition layer, comprising two linear layers, follows the sequence self-attention. This transition layer first expands the dimension and then projects it back to the original channel size. The resultant sequence representation is subsequently transformed into a pair representation through an outer product mean (OPM) block. The 2-D representation from the OPM block undergoes a sequential progression through a series of blocks, comprising (1) a triangle multiplicative update block using outgoing edges, (2) a triangle multiplicative update block using incoming edges, (3) a triangle self-attention block around the starting node, (4) a triangle self-attention block around the ending node, and (5) a pair transition block. It is important to highlight that the sequence and pair blocks are stacked residually for efficient and stable training.

The attention layer is configured with 8 heads, with each head having a channel size of 16. In the sequence transition layer, the expand factor is set to 2, meaning that the sequential representation is initially expanded to twice its original dimension. Within the OPM Module, the sequential representation is first projected into two representations with a channel size of 12, followed by the outer product operation to obtain a pair representation with a channel size of 12×12=144. This 144-D pair representation is then projected down to 64. For subsequent modules applied to this pair representation, the number of heads and the channel size of each head are also set to 8 and 16, respectively.

### Asymmetric modeling of TCR and peptide representations

The Joint Embedder and subsequent transformer block described above play crucial roles in integrating information across both TCR and peptide representations. Following this, the joint representations are spitted into TCR and peptide representations, each directed into separate branches of transformer blocks. This establishes an asymmetric deep learning structure for TCRs and peptides, characterized by the absence of parameters sharing between the two transformer blocks and further emphasized by a distinct layer configuration, i.e., N_1_=2 layers for TCRs and N_2_=1 layer for peptides (**Figure 1A**).

The asymmetry of the network is designed based on the empirical observation that the number of unique TCRs significantly exceeds that of unique peptides in the training set. Although alternative strategies, such as employing different hidden dimensions, are also plausible for achieving asymmetry, we opt for simplicity by differing in the block number and found that the latter design achieves slightly better performance compared to other selections.

### Training of TCR-peptide interaction model with triplet loss function design

Following the asymmetric modeling of TCR and peptide representations, the output TCR and peptide sequence representations are concatenated and go through an average pooling operation along the sequence axis, resulting a fixed-length vector. The final scores, which can quantify the interactions between the input TCR and peptide sequences, are then obtained through the final MLP module, which consists of 2 hidden layers with 126 and 64 neurons, respectively. (**Figure 1A**).

We have formulated a loss function based on the triplet loss for the training of TCRfinder. In each training step, a triplet of sequences is randomly sampled from the training set. This triplet consists of a peptide sequence (considered as the anchor), a TCR sequence that positively interacts with the peptide (denoted as *TCRp*), and a negative TCR sequence (*TCRn*).The TCRfinder model then computes the interaction scores between the anchor peptide and the two TCRs, i.e., *s*(*peptide, TCRp*) and *s*(*peptide, TCRn*) respectively, which are the target of interaction models. The loss function for each triplet is given by:

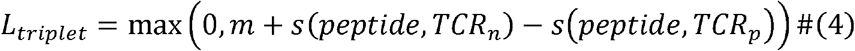

where *m* is the hyper-parameter that describes the margin and was set to 1.0 when training TCRfinder. Although the structure of the loss function in Eq. (3) resembles the previous triplet loss network protocol^34,35^, the learning in those triplet loss networks utilizes a given form of distance between samples, whereas in TCRfinder, the interacting score between difference sequences is unknown and trained as the target of the deep learning models.

In addition, inspired by the loss function utilized in pMTnet^12^, we introduce a new regularization term to Eq. (3) to regulate the scales of the scores. The ultimate form of the loss function is expressed as:

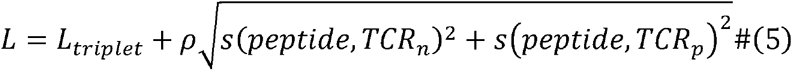

where *ρ* denotes the regularization factor and sets to 0.03 in the training of TCRfinder.

The training of TCRfinder’s prediction model involves the utilization of the Adam optimization method and the loss function defined in Eq (5), implemented with PyTorch. In each sampling step, one interacting and 10 non-interacting TCR sequences are randomly selected for a given peptide sequence. This sampling strategy has been observed to yield slightly improved performance compared to sampling across all possible pairs. The rationale behind this approach is the uneven distribution of interacting TCRs for each peptide. Directly sampling over all pairs may lead to overfitting on peptides with a larger number of interacting TCRs, and the adopted strategy aims to mitigate such potential biases.

The loss function for this specific sampling strategy is calculated as the average of ten triplet losses in a sampling step. In our formulation, we treat the sampling of one peptide and its interacting TCRs (including 10 negative TCRs) as a single sample, and one batch contains a fix number of 16 samples. Gradient accumulation has been utilized to reduce GPU memory consumption. Each of the 5 TCRfinder sub models has been trained for 10,000 batches and the early stopping mechanism is implemented to identify and select the optimal models.

### Alternative model trained for TCR-based neoantigen screening

To address the issue of TCR-based neoantigen screening, we have trained a separate model for recognizing mutant neoantigens from wildtype peptides under the same TCRs. For this, we have made straightforward adjustments to the sampling and loss function in the training process of the TCRfinder model, to enable peptide ranking given a TCR on the same dataset.

Each sample in the training set contains one TCR, one interacting peptide, and 10 non-interacting peptides randomly selected from the database. Consequently, the loss function for peptide ranking is formulated as:

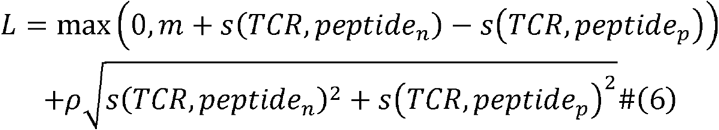

where *s*(*TCR,peptides*_*p*_) and *s*(*TCR,peptides*_*n*_) are the predicted scores between the TCR and interacting and non-interacting peptides, respectively. We trained a total of 5 models over 5,000 batches, implementing an early stopping mechanism to identify and select optimal models. The final score is the average score of the 5 models.

## DATA AVAILABILITY

The datasets collected and used in this work are available at https://zhanggroup.org/TCRfinder/dataset. We show structures of 6AMU obtained by four-digit accession codes in the PDB repository (https://www.rcsb.org/). TCR and peptide pairs for training were collected from VDJdb (https://vdjdb.cdr3.net/). The TCR and peptide sequences for training language models were downloaded from TCRdb (https://guolab.wchscu.cn/TCRdb//#/download) and IEDB (https://www.iedb.org/) respectively.

## CODE AVAILABILITY

The online server and standalone package of TCRfinder are freely available at https://zhanggroup.org/TCRfinder and https://zhanggroup.org/TCRfinder/download, respectively. Data were analyzed using Numpy v.1.20.3 (https://github.com/numpy/numpy), SciPy v.1.7.1 (https://www.scipy.org/), and Matplotlib v.3.4.3 (https://github.com/matplotlib/matplotlib). Structures were visualized by Pymol v.2.3.0 (https://github.com/schrodinger/pymol-open-source).

## ACKNOWLEDGMENTS

We thank Dr. Guideng Li and Chris Liu for insightful discussion.

## AUTHOR CONTRIBUTIONS

Y.Z. conceived the project and designed the experiments; Y.L. developed methods and designed and performed experiments; C.Z. participated in discussion and helped design the experiments; X.Z. constructed the online server. Y.L. wrote the initial manuscript; all authors proofread and approved the final manuscript.

