## Supporting Information for "TCRfinder: Improved TCR virtual screening for novel antigenic peptides with tailored language models"

### Supporting Texts

**Text S1: Dataset collections for Language model training and validation.**

We collected two types of datasets for LM development and TCR virtual screening experiments. For LM development, we collect all downloadable TCR sequences from TCRdb[1], comprising a total of 7,259,306 TCR β-chain CDR3 sequences, to train the TCR LM. The validation set for the TCR LM contains 884 TCRs included in VDJdb[2], which have the maximum sequence identity of 80% to the training TCRs.

In the case of the peptide language model, we collect all unique linear peptides from IEDB[3] and randomly select 1000 peptides for evaluation, with the remainder allocated for training. We also maintained a maximum sequence identity cutoff of 80% between the evaluation and training sets for peptide LM, and eventually the number of the training peptides is 1,500,786.

**Text S2: Dataset collections for TCR-peptide model construction and TCR and antigen screening.**

The training set for TCR-peptide interaction prediction was derived from VDJdb with following filters applied (1) TCR Species should be 'Human'; (2) Gene (chain) should be 'TRB'; (3) MHC Class should be 'MHCI'; and (4) Minimal confidence score should be above 1. As a result, a total of 286 unique peptides along with their corresponding interacting TCRs were extracted. 58 peptides (~20% of all training peptides) were randomly selected as validation set. The final pipeline is the average ensemble of 5 models trained with different parameter initializations and different training subsets with ~80% sampled data. Note that we also included all other TCR-peptide samples with confidence score = 0 during the training, as a simple data enrichment strategy.

For TCR virtual screening, since the aim of this study is to improve the TCR virtual screening task, especially with unseen peptides, the design of benchmark datasets is in the peptide level. A total of 38 unseen peptides were collected from the VDJdb database of curated TCR sequences with known antigen specificities[2], as the independent dataset. The verification T-cell specificity score was set to 3, meaning that the collected TCR-peptide interactions have extensive verification or structural data. Here, 'unseen' means that the peptide was never showed up in any training sets of the TCRfinder or control methods. Each unseen peptide contains 1 or at most 239 interacting TCRβ CDR3 sequences. Another 5,000 TCRβ CDR3 sequences were randomly selected as background sequences from TCRdb[1] database, where the goal of the task is to identify the unseen peptide specific TCRs against 5,000 background sequences.

For neoantigen screening, we removed any pairs in the VDJdb database that share the same TCR or peptide sequences with the 32 test cases. Considering the aim of this task is to improve antigen virtual screening, the remaining pairs were grouped by TCRs, resulting in 33,648 unique TCRs with confidence score $\geq$ 0. Similarly, we trained five models, each of which was validated on approximately 670 randomly selected TCRs, with the remaining TCRs and their interacting peptides used for training.

### Supporting Figures

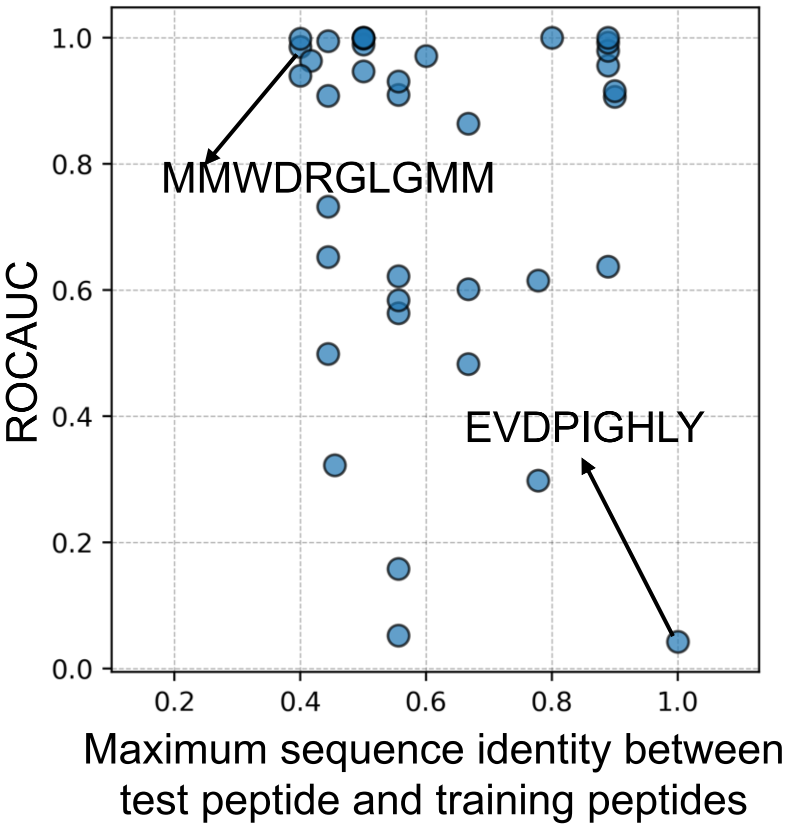


**Figure S1. Relationship between ROCAUC and maximum sequence identity among peptides in the test and training sets.**

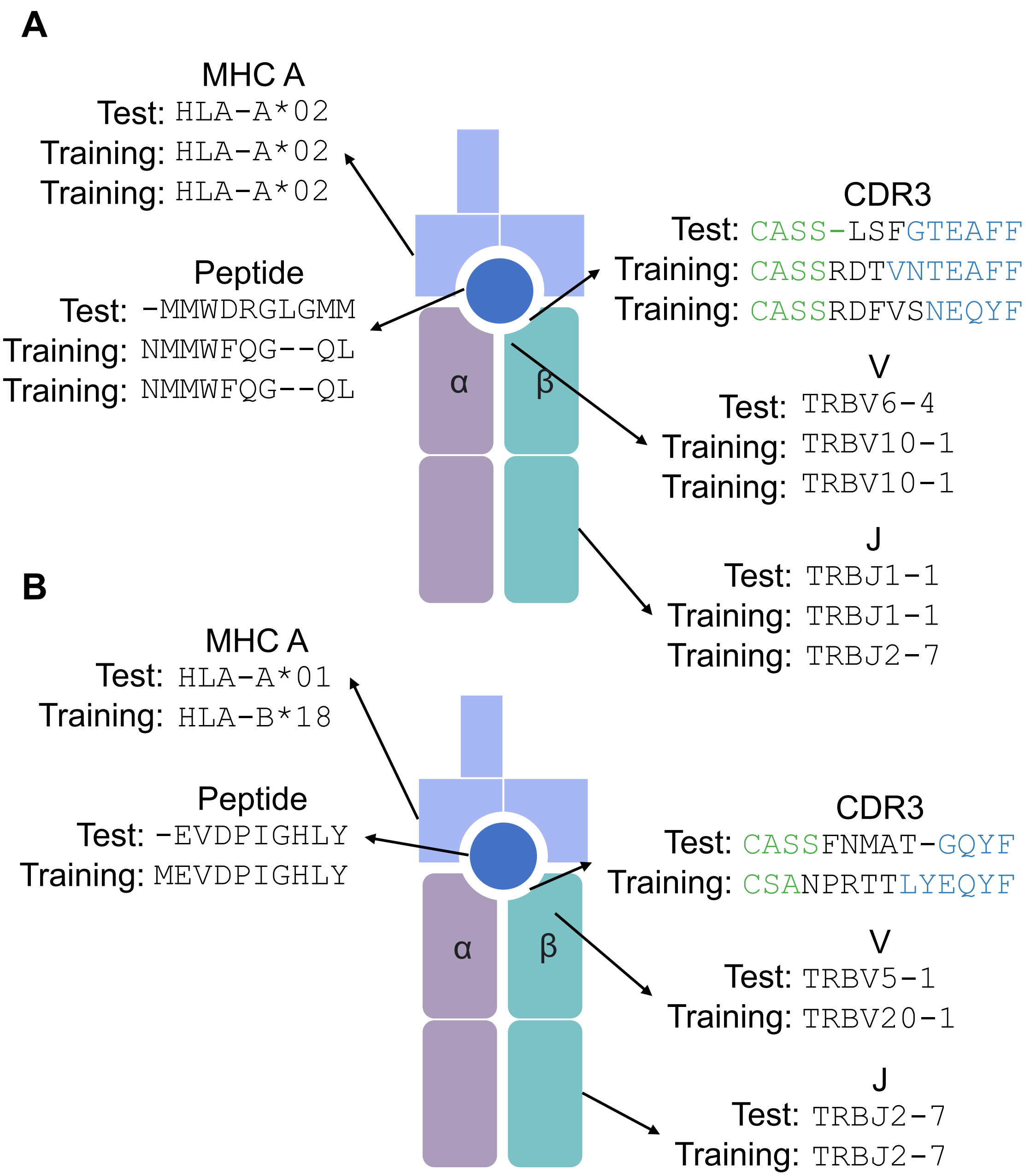


**Figure S2 Comparison of test TCR-pMHC pairs (labeled with 'Test') with the one having the highest peptide sequence identity in the training set (labeled as 'Training').** (A) Comparison for case peptide 'MMWDRGLGMM'. (B) Comparison for case peptide 'EVDPIGHLY'.

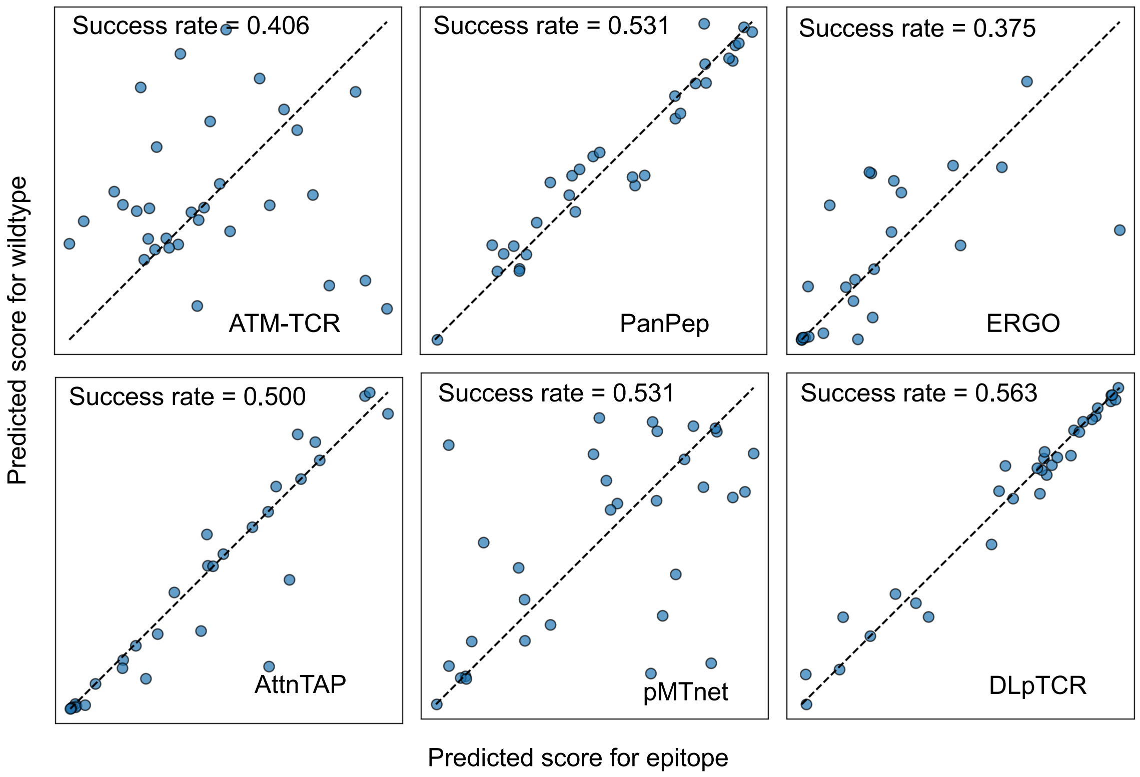


**Figure S3. Success rates of control method on the dataset with peptides derived from neoantigens and their corresponding wild types.**

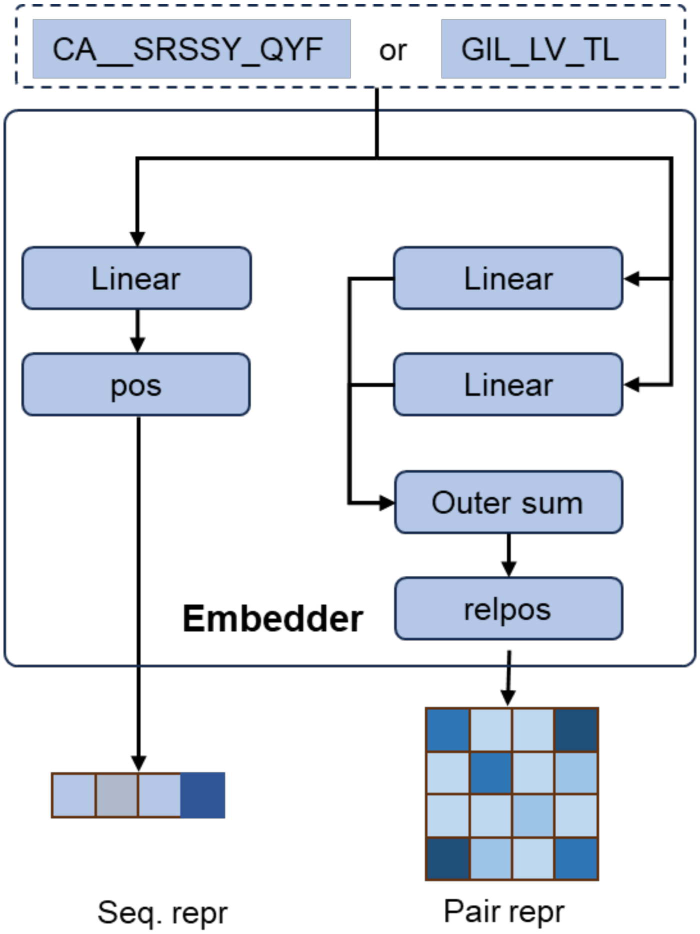


**Figure S4. Detailed flowchart illustrating the Embedder Module of TCR and peptide language models.**
